# Early functional connectivity alterations in contralesional motor networks influence outcome after severe stroke

**DOI:** 10.1101/2023.03.08.531668

**Authors:** Hanna Braaß, Lily Gutgesell, Winifried Backhaus, Focko L. Higgen, Fanny Quandt, Chi-un Choe, Christian Gerloff, Robert Schulz

## Abstract

Connectivity studies have significantly extended the knowledge on motor network alterations after stroke. Compared to interhemispheric or ipsilesional networks, changes in the contralesional hemisphere are poorly understood. Data obtained in the acute stage after stroke and in severely impaired patients are remarkably limited. This study aimed to investigate early functional connectivity changes of the contralesional parieto-frontal motor network and their relevance for the functional outcome after severe motor stroke. Resting-state functional imaging data were acquired in 19 patients within the first two weeks after severe stroke. Nineteen healthy participants served as a control group. Functional connectivity was calculated from five key motor areas of the parieto-frontal network on the contralesional hemisphere as seed regions and compared between the groups. Connections exhibiting stroke-related alterations were correlated with clinical follow-up data obtained after 3 to 6 months. The main finding was an increase in coupling strength between the contralesional supplementary motor area and the sensorimotor cortex. This increase was linked to persistent clinical deficits at follow-up. Thus, an upregulation in contralesional motor network connectivity might be an early pattern in severely impaired stroke patients. It might carry relevant information regarding the outcome which adds to the current concepts of brain network alterations and recovery processes after severe stroke.

## Introduction

Magnetic resonance imaging (MRI) based connectivity analyses have significantly contributed to our present understanding of how functional brain networks are affected by acute stroke lesions and how these changes are associated with residual motor function and recovery processes ^1^. Most evidence has been accumulated for networks connecting sensorimotor cortices of both hemispheres with preserved or increased interhemispheric connectivity being positively related to favorable outcomes ^2–5^. Compared to interhemispheric connectivity, there are few data to support the view that alterations in intrahemispheric coupling profiles within the ipsilesional ^6–9^ and especially the contralesional hemisphere ^10–13^ might explain inter-subject variability in motor functions or recovery after stroke. MRI data from clinical cohorts investigating the contralesional coupling changes obtained in the acute to the early subacute stage are remarkably scarce but functional ^14–16^ and structural MRI imaging ^17–19^, and non-invasive brain stimulation ^20–22^ have convergingly indicated that the sensorimotor parieto-frontal network of the contralesional hemisphere is involved in recovery after stroke, with greater importance in more severely impaired patients. The study aimed to investigate early functional network changes with a focus on key areas of the contralesional parieto-frontal motor network and their relevance for the functional outcome after severe motor stroke. Structural and resting-state functional MRI data, obtained within the first two weeks after severe stroke, were re-analyzed from a previously published prospective cohort study of severely impaired acute stroke patients and integrated with clinical follow-up data of the late subacute stage of recovery after 3 to 6 months ^6^. A seed-based approach was used to assess contralesional resting-state functional connectivity (FC) involving five key motor areas, that are the primary motor cortex (M1), ventral premotor cortex (PMV), supplementary motor area (SMA), and anterior and caudal intraparietal sulcus (AIPS, CIPS). We hypothesized that stroke association between FC estimates and subsequent recovery.

## Results

### Demographics and clinical characteristics

Nineteen patients (12 females and 7 males, all right-handed, aged 73.8 ± 5.8 years) and 19 healthy controls (12 females and 7 males, all right-handed, aged 75.3 ± 7.5 years) were included in the analysis. A topographic map of the distribution of stroke lesions is shown in Figure 1. Clinical characteristics are given in Supplementary Table 1. Early clinical examination was conducted on average on day 7 (mode day 5, range 3–13) after stroke. Late sub-acute stage follow-up data (LSA) were derived from clinical examination after 128 days on average (mode 89, range 86–217). Linear mixed-effects models evidenced significant functional improvements over time in ‘early rehabilitation’ Barthel Index (BI) ^23^, Fugl Meyer Assessment of the upper extremity (UEFM), modified Rankin Scale (MRS), and National Institutes of Health Stroke Scale (NIHSS) (all *P*<0.001, details are shown in ^6^).

**Figure 1.**
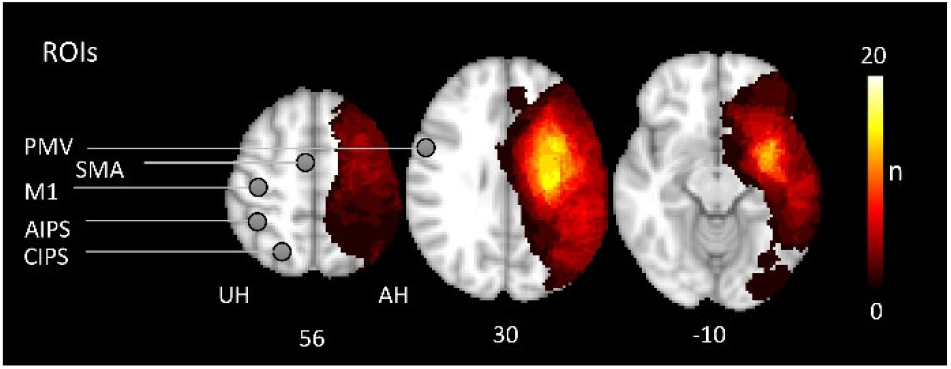
Stroke lesions and motor network regions of interest (ROI) All masks of stroke lesions are displayed in the left hemisphere (affected hemisphere, AH), overlaying a T1-weighted template in MNI space (z-coordinates below each slice). The color intensity indicated the number of subjects of whom lesion voxels lay within the colored region. Contralesional motor ROIs (M1, PMV, SMA, AIPS and CIPS) are displayed on the unaffected hemisphere (UH).

### Alterations in contralesional functional connectivity after stroke

Seed-based whole brain analyses revealed FC alterations between contralesional seed areas M1, PMV, SMA, and CIPS and multiple brain regions of both hemispheres compared to healthy controls in the acute phase after stroke. No significant differences were found for AIPS. Table 1 shows the group differences for contralesional FC between stroke patients and controls and Figure 2 illustrates the topography of contralesional clusters exhibiting FC alterations after stroke. For M1, we detected an increase in FC with precuneus, cingulate gyrus, SMA, and precentral gyrus (region of the dorsal premotor cortex). SMA was found to be more strongly connected to the sensorimotor cortex (SMC), more precisely, the peak coordinate of the SMC cluster is located in the sulcus centralis between contralesional precentral and postcentral gyrus. Increases in FC were also detected for SMA and frontal pole, middle frontal gyrus (MFG), lateral occipital cortex, and precuneus. PMV exhibited stronger FC with frontal pole and MFG, and CIPS was more strongly coupled with a cluster localized in precuneus and cingulate cortex.

**Figure 2.**
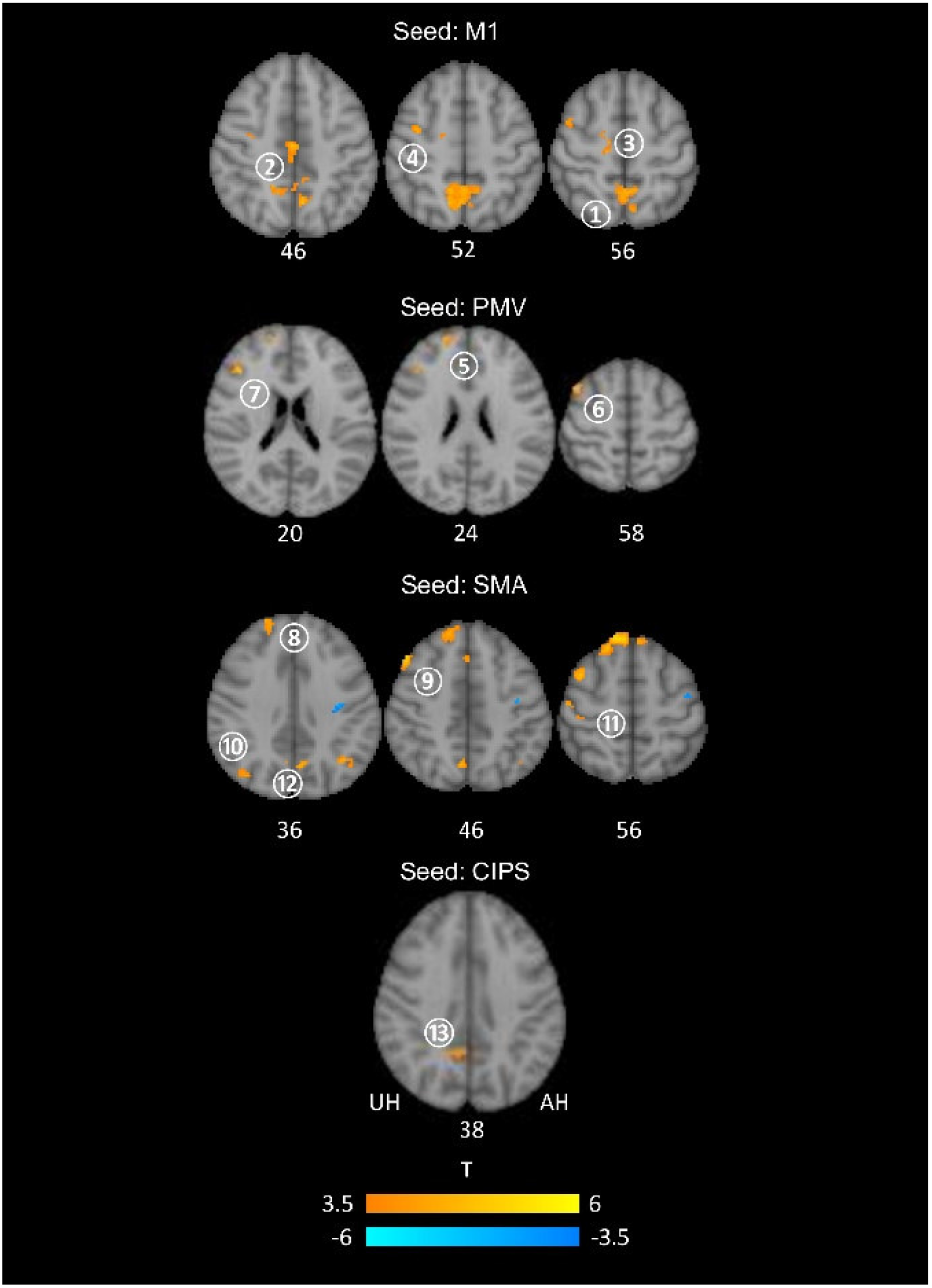
FC changes after stroke from contralesional seed regions compared to healthy controls. Clusters exhibiting significant changes in FC for M1, PMV, SMA and CIPS as contralesional seed regions are superimposed on a standard T1 image in MNI space. T values are color-coded with orange yellow indicating an increase in FC in stroke patients, and blue indicating a decrease in FC. The numbers of the unaffected (contralesional) hemisphere refer to the cluster IDs used in table 1. Z-values in MNI space are given below each axial slice. AH affected hemisphere, UH unaffected hemisphere. For details of contralesional clusters, please see table 1.

**Table 1.** FC changes in the contralesional hemisphere compared to health controls. Clusters exhibiting significant increases in patients compared to controls are given with region information, MNI coordinates, cluster sizes, FDR-corrected P and T values. Results are derived from a cluster size threshold of P<0.05 (FDRcorrected) and a voxel threshold of P<0.001 (uncorrected). The Harvard Oxford atlas was used for the region classification.

| Seed | Region | Cluster | MNI Coord. | | | Cluster size | $P_{FDR}$ | T |
| --- | --- | --- | --- | --- | --- | --- | --- | --- |
|  |  |  | x | y | z |  |  |  |
| M1 | Precuneus | 1 | 0 | -40 | 66 | 590 | <0.0001 | 5.5 |
|  | Cingulate Gyrus | 2 | 2 | -16 | 42 | 146 | <0.001 | 4.86 |
|  | SMA | 3 | 10 | -14 | 60 | 55 | 0.026 | 4.73 |
|  | Precentral Gyrus | 4 | 36 | -6 | 52 | 51 | 0.029 | 4.5 |
| PMV | Frontal Pole | 5 | 20 | 56 | 24 | 105 | 0.008 | 4.51 |
|  | Middle Frontal Gyrus (MFG) | 6 | 42 | 12 | 60 | 58 | 0.03 | 5.10 |
|  | MFG | 7 | 42 | 30 | 20 | 56 | 0.033 | 5.21 |
| SMA | Frontal Pole | 8 | 12 | 44 | 52 | 363 | <0.0001 | 5.58 |
|  | MFG | 9 | 48 | 26 | 46 | 222 | <0.0001 | 5.27 |
|  | Lateral Occipital Cortex | 10 | 34 | -60 | 28 | 75 | 0.013 | 4.31 |
|  | Postcentral/Precentral Gyrus (SMC) | 11 | 48 | -16 | 58 | 55 | 0.025 | 4.88 |
|  | Precuneus | 12 | 6 | -64 | 46 | 51 | 0.031 | 4.58 |
| CIPS | Precuneus/Cingulate Gyrus | 13 | 8 | -50 | 34 | 74 | 0.045 | 4.54 |

### Associations between altered contralesional functional connectivity and future persistent deficits after stroke

Linear models were fitted to relate FC values in the acute phase to clinical outcome after 3-6 months, operationalized by means of BI, UEFM, MRS and NIHSS (Table 2). The main finding was a consistent association of the FC between contralesional SMA and SMC and clinical outcome and upper limb function. An early increase in FC was linked to lower scores in BI (*P*=0.002), in UEFM (*P*=0.018) and higher scores in MRS (*P*=0.024) at follow-up. For NIHSS, we observed a compatible statistical trend (*P*=0.067). Importantly, the association of FC values obtained early after stroke and functional outcome at follow-up was independent from the initial deficit. Notably, there was no correlation between FC values and initial deficits. The addition of SMA-SMC FC to the initial behavioral scores increased the explained variance by 30% in future BI, 24.5% in UEFM, and 22% in MRS, respectively (see Table 3 and Figure 3 for statistical model results). Further, FC-outcome associations with the same direction were detected for M1-SMA FC and MRS (*P*=0.038), PMV-MFG FC and MRS (*P*=0.038), and CIPS-Precuneus/Cingulate Gyrus FC and NIHSS (*P*=0.047) (Table 3).

**Table 2.**
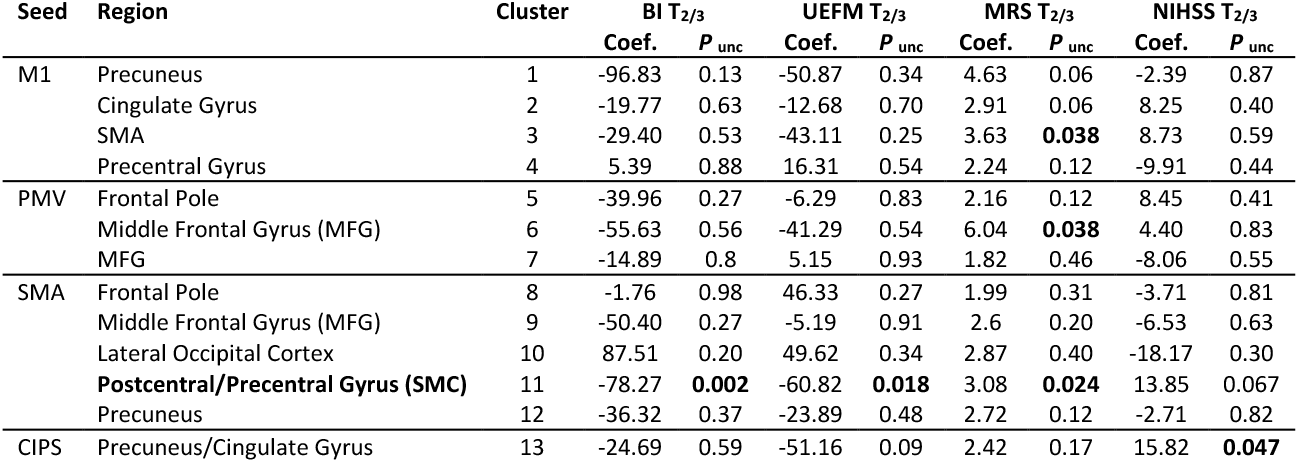
Association of contralesional functional connectivity and future persistent deficits. Coefficients are given incl. their *P*-values (uncorrected within models) for contralesional FC (obtained at timepoint T_1_) as the main predictor. Results are derived from independent models for the four outcome scores and follow-up timepoint T_2/3_. Significant findings are highlighted in bold. See table 3 for further model details for SMA-SMC FC (cluster 11).

**Table 3.**
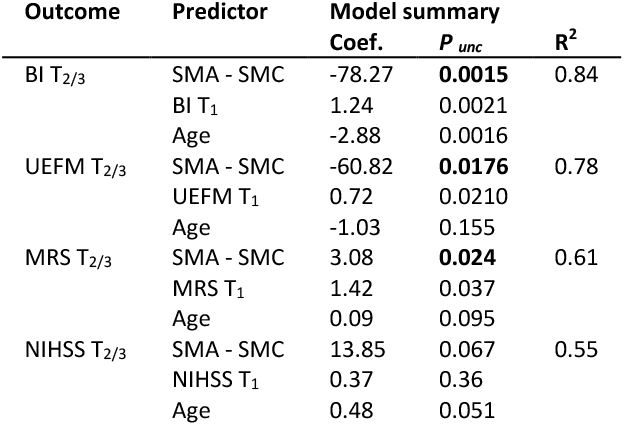
Association of contralesional SMA-SMC FC and future persistent deficits. Coefficients are given incl. their *P*-values (uncorrected within models) for contralesional SMA-SMC FC (cluster 10, obtained at timepoint T_1_) as the main predictor for the four outcome scores and follow-up timepoint T_2/3_. FC-outcome relationships are adjusted for the influence of the initial deficit and age. R^2^ shows multiple R^2^. Significant findings are highlighted in bold.

| Outcome | Predictor | Model summary |  | R<sup>2</sup> |
| --- | --- | --- | --- | --- |
|  |  | Coef. | P<sub>unc</sub> |  |
| BI T<sub>2/3</sub> | SMA - SMC | -78.27 | **0.0015** | 0.84 |
|  | BI T<sub>1</sub> | 1.24 | 0.0021 |  |
|  | Age | -2.88 | 0.0016 |  |
| UEFM T<sub>2/3</sub> | SMA - SMC | -60.82 | **0.0176** | 0.78 |
|  | UEFM T<sub>1</sub> | 0.72 | 0.0210 |  |
|  | Age | -1.03 | 0.155 |  |
| MRS T<sub>2/3</sub> | SMA - SMC | 3.08 | **0.024** | 0.61 |
|  | MRS T<sub>1</sub> | 1.42 | 0.037 |  |
|  | Age | 0.09 | 0.095 |  |
| NIHSS T<sub>2/3</sub> | SMA - SMC | 13.85 | 0.067 | 0.55 |
|  | NIHSS T<sub>1</sub> | 0.37 | 0.36 |  |
|  | Age | 0.48 | 0.051 |  |

**Figure 3.**
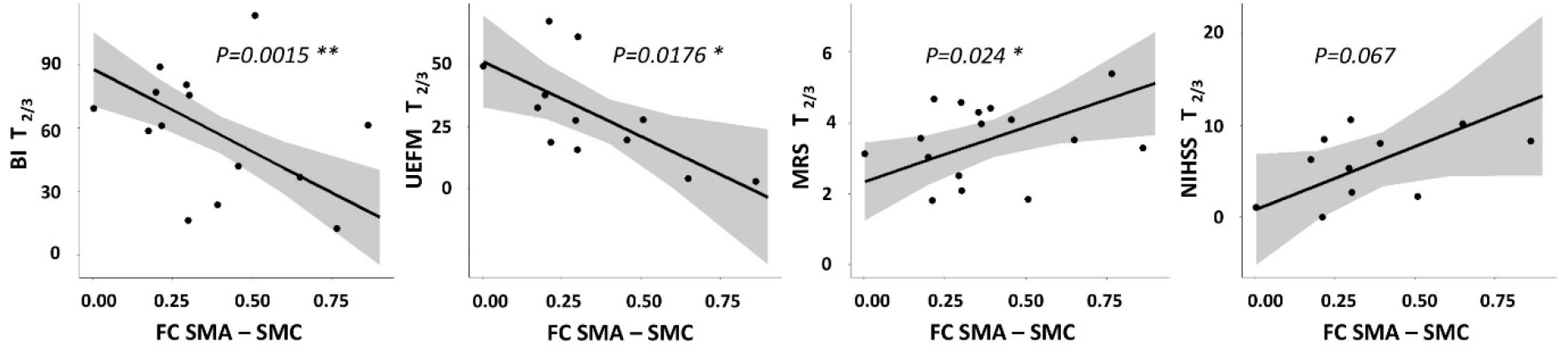
Influence of contralesional FC on future persistent deficits after stroke. Effect plots are shown for contralesional SMA-SMC FC contributing to the explanation of variability in follow-up BI, UEFM, MRS and NIHSS in severe stroke patients. There was a significant association between FC SMA-SMC at T1 and BI, UEFM and MRS at T_2/3_ with higher FC values early after stroke found in patients which are likely to show more severe persistent deficits in follow-up. *P* of FC SMA-SMC as the predictors of interests (within-model) is given (uncorrected).

## Discussion

In patients with severe stroke at initial presentation, we found an increase in contralesional functional connectivity between SMA and sensorimotor cortex (SMC). The increase in FC was negatively associated with the extent of persistent deficits at follow-up after 3-6 months, independently of the initial deficits and age.

The present findings of an early increase in contralesional FC is well in line with previous work showing an upregulation in activity in contralesional motor areas including SMA and SMC, particularly in more severely impaired patients with poor motor outcome ^14,15^. For instance, one study in 11 acute stroke patients found a global reduction of task-related activity acutely after stroke followed by increases in ipsilesional and contralesional motor areas within the first 10 days ^24^. Such increases in contralesional activation might persist over time even in well-recovered patients ^25,26^. In contrast, other studies found downregulated activity in contralesional SMA in the chronic stage of recovery ^27,28^. Inter-study variability, e.g., in the level of impairment and time points, leads to a rather complex picture of time- and recovery-dependent changes in brain activation ^16^. In addition to local activation, stroke also leads to widespread changes in inter-regional connectivity. In fact, most evidence is available for interhemispheric connections and intrahemispheric connections of the ipsilesional hemisphere ^1,6–9^. Data on contralesional coupling dynamics and their importance for recovery are remarkably limited. The situation is further complicated by a large variability in clinical cohorts, methods and time points investigated. For instance, in well-recovered patients and the chronic stage of recovery, affected hand movements have been linked to an impaired task-related inhibitory information flow from contralesional SMA to contralesional M1 ^10^. In the acute setting, another study observed a reduced inhibitory information flow from SMA to contralesional M1 ^11^. An analysis in 81 patients, measured within two weeks after ischemic or hemorrhagic stroke, has found a decrease in resting-state FC between contralesional SMA and M1 ^12^. However, none of these studies have reported any coupling-outcome associations. In contrast, one study in severely impaired chronic stroke patients has revealed increased connectivity in the contralesional sensorimotor cortex within the sensorimotor network, when compared to less impaired patients. The level of upregulated contralesional coupling was negatively correlated with the performance of the paretic hand ^13^. The present results add to these data by indicating that the upregulation in contralesional connectivity might be an early and specific pattern in severely impaired stroke patients which might also carry, as evidenced for SMA-SMC, relevant information regarding future persistent deficits. As this association was independent of the initial deficit, we argue that the increase in FC might parallel the attempt of the lesioned brain to recruit additional motor cortices of the contralesional hemisphere to support motor functions and facilitate recovery. However, this attempt appears to be largely futile in the end. To what extent an additional contralesional upregulation by means of non-invasive brain stimulation ^29^ might help to promote recovery has to be investigated systematically and prospectively in large, well-characterized clinical cohorts. Thereby, models of interhemispheric inhibition suggesting potentially non-beneficial effects in the ipsilesional hemisphere should not stand against these considerations as more recent data have just begun to call those concepts into question ^30^. Apart from SMA-SMC, FC increases were also present in the contralesional precuneus and cingulate cortices, particularly seeding from M1 and CIPS. These alterations are well in line with previous data obtained in severely impaired acute stroke patients, which also showed an increase in specific states of dynamic FCs between the precuneus and frontal brain regions ^31^. Chronic stroke patients comparably exhibited a stronger coupling between medial regions of the posterior default mode network and the frontoparietal network ^32^. The absence of significant associations between these alterations and subsequent motor outcome in the present cohort might be explained by the actual focus on the motor domain. Poststroke cognitive impairment and affective symptoms have been associated with FC alterations in default mode network regions ^33,34^.

There are some critical limitations worth noting. First, the authors fully recognize the small sample size, which is likely to reduce the sensitivity and specificity of the present findings. Whole-brain results are corrected for multiplicity in line with the methodological standard. However, subsequent correlations were not corrected for multiple testing, which biases the results towards higher sensitivity at lower specificity. Though, the consistency of SMA-SMC association with multiple outcome scores suggests a specific and valid finding. Nevertheless, these results remain exploratory in nature; upcoming studies will have to verify the findings. Second, as seed areas, we selected five key areas of the contralesional parieto-frontal motor network. Whether the FC changes might be mediated by hidden, unmodeled nodes remains unclear. Third, seed-based FC analyses were computed across the whole brain. However, following our a-priori hypothesis, further statistical analyses were restricted to the contralesional hemisphere as direct lesion effects might be very difficult to quantify and critically influence FC estimates. However, the correction for multiplicity across the brain is still influenced by the ipsilesional hemisphere.

## Materials and methods

### Cohort and clinical data

The present analyses are based on clinical and imaging data of a previously published prospective cohort study comprising 30 more severely impaired acute stroke patients admitted to the University Medical Center Hamburg-Eppendorf from October 2017 to February 2020 ^6^. The study was approved by the local ethical commission (PV5442). Acute stroke patients (3 -14 days after the incident) were included according to the following criteria: first-ever ischemic stroke causing a severe motor deficit involving hand function, modified Rankin Scale (MRS) > 3 or Barthel index (BI) ≤ 30 or ‘early rehabilitation’ Barthel Index < 30 and age ≥ 18 years. Exclusion criteria were pre-existing clinically silent brain lesions > 1 cm^3^ or pre-existing motor deficits, contraindications for MRI, relevant psychiatric diseases, drug abuse or pregnancy. A flowchart of study inclusion is given in the original report ^6^. The participants provided informed consent themselves or via a legal guardian, following the ethical Declaration of Helsinki. Acute stroke patients underwent structural and functional resting-state MRI in the first days after the event as time point T_1_ (days 3-14). Follow-up time point T_2_ was defined in the late subacute stage of recovery ^35^ after three months, or, in patients in which clinical data for this time point was not available, after six months. Standardized tests at time point T_1_ and T_2_ included the BI, the Fugl Meyer Assessment of the upper extremity (UEFM), the MRS, and the National Institutes of Health Stroke Scale (NIHSS). Nineteen Patients were matched with 19 healthy control participants according to age and sex. All patients and controls were right-handed.

### Brain Imaging – Data acquisition

A 3T Skyra MRI scanner (Siemens Healthineers, Erlangen, Germany) equipped with a 32-channel head coil were used to acquire multimodal imaging data, including structural high-resolution T1-weighted images and functional resting-state images. For the T1-weighted sequence, a 3-dimensional magnetization-prepared rapid gradient echo (3D-MPRAGE) sequence was used with the following parameters: repetition time (TR)=2500ms, echo time (TE)=2.12ms, flip angle 9°, 256 coronal slices with a voxel size of 0.8×0.8×0.9mm^3^, field of view (FOV)=240 mm. The resting-state parameters for blood oxygenation level-dependent (BOLD) contrasts were FOV=260mm, TR=2 s, TE=30ms, a 72×72×32 matrix, voxel size 3×3×3mm^3^, flip angle 90°, and 210 images. Before the resting-state scans, the participants were asked to focus on a black cross located behind the scanner, which could be viewed via a mirror. For analyses, all resting-state and T1-weighted images with right-sided stroke lesions were flipped to the left hemisphere. This hemispheric flip was performed in the respective matched controls to account for the distribution of stroke lesions to the dominant and non-dominant hemispheres in line with the original report ^6^.

### Brain Imaging – Image analysis

The resting-state images and T1-weighted images were preprocessed using the CONN toolbox v20.b, an SPM12-based toolbox. For image preprocessing the default-pipeline for volume-based analysis was used with the following steps and parameters: removal of the initial 10 scans and application of a 6 mm full width at half maximum (FWHM) Gaussian kernel for smoothing of the functional images, normalization and denoising using the anatomical component-based (i.e., white matter, CSF, realignment, and scrubbing) noise correction procedure (*a*CompCor) for BOLD-based functional MRI ^36^. Global signal regression was not included in the analysis to avoid potential false anti-correlations ^37^. A temporal band-pass filter between 0.008 Hz and 0.1 Hz was applied to focus on slow-frequency fluctuations while minimizing the influence of physiological, head motion, and other noise sources ^38^.

### Seed-based analysis

In a first level analysis the interregional functional connectivity (FC) was conducted following a whole-brain approach (seed-based analysis) using spherical seeds (radius of 5mm) for five key areas of the parieto-frontal motor network of the contralesional hemisphere with published MNI coordinates ^9^: the primary motor cortex M1 (38, -22, 54), the supplementary motor area SMA (6, -4, 57), the ventral premotor cortex PMV (54, 6, 32), the anterior intraparietal sulcus AIPS (38, -43, 52) and the caudal intraparietal sulcus CIPS (21, -64, 55). Voxel-wise FC values were calculated between these seeds and all other voxels based on Fisher-transformed bivariate cross-correlation coefficients. For group-wise second level, cluster-based analyses were performed for comparison between patients and controls [1 -1] with a cluster size threshold of *P*<0.05 (FDR-corrected) and a voxel threshold of p<0.001 (uncorrected). The results were used to identify the peak coordinates of the different clusters and create spherical regions of interest (ROI, radius of 5mm) around these peak coordinates. The ROIs were used in ROI-to-ROI analyses to obtain individual Fisher-transformed bivariate correlation coefficients (FC) between each pair of seed and peak ROI for further correlational analyses.

### Statistical Analysis

Statistical analyses were performed in R (version 4.0.4). To assess functional improvement over time, linear-mixed effects models with repeated measures were fitted with TIME as the factor of interest and ID as random effect. If available, the three months follow-up time point was used, otherwise clinical data after six months were used in line with the original work ^6^. To assess FC-outcome relationships, individual linear models were constructed with follow-up BI, UEFM, MRS or NIHSS as dependent variable and cluster-wise FC value as the predictor of interest and the initial deficit at T_1_ (equivalent score) and AGE as covariates to adjust the target effects. Model results are presented by predictor coefficients with their significances and overall explained variance of the final models. Statistical significance was set to a *P*<0.05 (uncorrected).

## Author contributions

Study Design: R.S., W.B.

Data acquisition: W.B., F.H.,

Data analysis and statistical analysis: H.B., L.G., R.S.

Data interpretation: H.B., L.G., F.H., R.S., F.Q., C.U.C., C.G.

First Manuscript: H.B., L.G.

All authors reviewed the manuscript.

## Data availability

Data will be made available upon reasonable request, which includes submitting an analysis plan for a secondary project.

