## Supplementary material for "Early functional connectivity alterations in contralesional motor networks influence outcome after severe stroke"

| ID | Age | Sex | Side / Dom | **TL / MT** (TICI) | LVO | LV (ml) | Days C/I | NIHSS | | UEFM | | MRS | | BI | |
|---|---|---|---|---|---|---|---|---|---|---|---|---|---|---|---|
| | | | | | | | | Acute | LSA | Acute | LSA | Acute | LSA | Acute | LSA |
| 1 | 78 | female | left / d | no / no | none | 33.6 | 7/7 | 10 | 3 | 8 | 31 | 4 | 3 | 35 | 80 |
| 2 | 63 | male | left / d | no / no | M1 | 55.8 | 3/3 | 13 | 1 | 5 | 36 | 4 | 1 | 40 | 100 |
| 3 | 73 | female | left / d | yes / yes (2B) | M1 | 14.4 | 5/5 | 9 | 3 | 48 | 65 | 4 | 3 | 20 | 100 |
| 4 | 73 | female | right / n | yes / yes (2A) | M1 | 27.6 | 5/5 | 5 | 2 | 49 | 62 | 4 | 1 | 25 | 80 |
| 5 | 79 | female | right / n | yes / yes (2B) | M2 / A1 | 120.4 | 5/6 | 8 | 2* | 15 | 51* | 5 | 4 | 10 | 85* |
| 6 | 89 | female | right / n | yes / no | none | 2.6 | 4/4 | 7 | 3 | 5 | 39 | 5 | 3 | 40 | 55 |
| 7 | 71 | female | right / n | yes / yes (2A) | ACI/M1/A1 | 38.4 | 8/8 | 9 | - | 4 | 47* | 5 | 3* | 10 | 70* |
| 8 | 76 | male | right / n | yes / yes (3) | M1/A2 | 101.0 | 5/6 | 11 | - | 4 | - | 5 | 3† | 10 | - |
| 9 | 78 | male | right / n | yes / yes (3) | M2 | 178.1 | 4/4 | 17 | 3 | 5 | 15 | 5 | 4 | 10 | 65 |
| 10 | 85 | female | right / n | no / yes (2B) | M1 | 33.5 | 5/5 | 15 | 14 | 2 | 4 | 5 | 5 | 0 | 0 |
| 11 | 78 | male | left / d | no / no | M1 | 58.1 | 10/15 | 17 | - | 3 | - | 5 | 5*,† | 10 | 15*,† |
| 12 | 74 | male | left / d | yes / yes (2B) | M1 | 303.3 | 10/10 | 24 | - | 5 | 4* | 5 | 5 | 5 | 5 |
| 13 | 69 | male | left / d | yes / yes (2A) | M1 | 98.4 | 7/7 | 18 | - | 0 | - | 5 | - | 0 | - |
| 14 | 77 | female | right / n | yes / no | M1 | 286.7 | 7/7 | 11 | 10* | 4 | 4* | 4 | 4* | 20 | 45* |
| 15 | 67 | female | right / n | yes / no | none | 7.4 | 8/7 | 11 | 7 | 6 | 5* | 4 | 3 | 30 | 95 |
| 16 | 58 | female | left / d | no / no | ACI/M1 | 58.4 | 13/13 | 23 | - | 0 | - | 5 | - | 5 | - |
| 17 | 80 | female | left / d | no / no | M1 | 20.5 | 12/12 | 11 | 15* | - | 4* | 5 | 4* | 0 | 30* |
| 18 | 83 | female | left / d | yes / yes (3) | Carotis-T | 21.6 | 9/9 | 10 | - | 6 | - | 5 | - | 20 | - |
| 19 | 80 | male | right / n | yes / yes (0) | M1 | 108.4 | 7/7 | 16 | - | 6 | - | 5 | 6† | 10 | - |
| **Stroke** | 73.8 (5.8) | 7 male | 10 right / 9 d | 13 TL / 11 MT | mode: M1 | 82.5 (87.6) | mode: 5/7 | 12.89 (5.17) | 5.7 (5.0) | 9.7 (14.5) | 30.3 (23.3) | 4.7 (0.5) | 3.6 (1.4) | 15.8 (13.0) | 58.9 (34.8) |
| **Controls** | 75.3 (7.5) | 7 male | - | - | - | - | | - | - | - | - | - | - | - | - |

**Supp. Tab. 1| Patient characteristics at baseline and functional improvement over time.**
Baseline characteristics of all patients individually and averaged per group, for patients and controls. Mean values and standard deviation in brackets are given. Side lesion side. Dom Hemispherical dominance with d dominant hemisphere, n non-dominant hemisphere affected by the lesion. TL thrombolysis, MT mechanical thrombectomy, TICI: thrombolysis in cerebral infarction grading system, partial perfusion of the treated vessel is reached in grade 2B (mode). LVO large vessel occlusion. LV lesion volume. M1/M2/A1/ACI indicate the cerebral vessels occluded. Days C/I Exam time point of clinical examination (C) and imaging (I) in days after stroke. Recovery and initial scoring of four scores: NIHSS National Institute of Health Stroke Scale, UEFM Upper Extremity Fugl Meyer Assessment, MRS modified Rankin Scale, BI Barthel-Index. Time point of data collection in the acute (day 4-14) or late sub-acute stage (LSA), either three or six months (depicted by *) after stroke. † indicate follow-up values collected via phone.
